# Indole toxicity on removal of uremic toxin p-cresol, in-vitro study of *Thauera aminoaromatica* S2

**DOI:** 10.64898/2026.01.30.702922

**Authors:** Pei-Hsin Wang, Prakit Saingam, Rosita Rasyid, Bruce J.Godfrey, Jonathan Himmelfarb, Mari Karoliina Henriikka Winkler

## Abstract

Protein-bound uremic toxins, such as indoxyl sulfate and *p*-cresyl sulfate, are major contributors to chronic kidney disease (CKD) complications and are poorly removed by dialysis due to strong albumin binding. Targeting their gut-derived microbial precursors offers a promising strategy to reduce systemic toxin load. *Thauera aminoaromatica* S2 is known to anaerobically degrade *p*-cresol, but its response to indole and its potential as an orally administered microbial therapy remain poorly characterized. Here, we investigated the activity of *Thauera aminoaromatica* S2 under exposure to both *p*-cresol and indole in planktonic and hydrogel-encapsulated forms. Low indole levels (0.25 mM) enhanced planktonic growth in the presence of 2 mM *p*-cresol, whereas co exposure inhibited p-cresol degradation in hydrogel systems, likely due to restricted diffusion and elevated local indole concentrations. Nonetheless, encapsulation enabled tolerance to conditions (2 mM *p*-cresol + 0.5 mM indole) that abolished planktonic growth, suggesting microenvironmental protection. Incorporation of activated carbon into the hydrogel restored *p*-cresol removal despite indole exposure, likely through localized indole sequestration. These results highlight the potential of combining encapsulation with adsorptive additives to stabilize microbial function and support the development of microbial therapies aimed at mitigating uremic toxin precursors in CKD.

## Introduction

Patients with chronic kidney disease (CKD) gradually lose kidney function, which leads to risk of complications and mortality (Narayanan and Setia 2019). Uremic toxins accumulate in the blood when kidney function declines and contributes to CKD progression and systemic complications (Fujii, Goto, and Fukagawa 2018; Kilmer 2010). These toxins are broadly classified into three categories: water-soluble low–molecular-weight solutes, middle molecules, and protein-bound uremic toxins (PBUTs) (Duranton et al. 2012). Among them, PBUTs are the most challenging to eliminate because their strong binding to plasma proteins severely limits removal by conventional dialysis (Liu, Tomino, and Lu 2018). *P*-cresol and indole are protein bound uremic toxins formed from the intestinal microbial metabolism of the aromatic amino acids, tyrosine and tryptophan, respectively. (Evenepoel et al. 2009). Indole and *p*-cresol are sulfonated to indoxyl sulfate and *p*-cresyl sulfate in the liver and eventually removed by tubular secretion in the kidney (Gryp et al. 2017; Leong and Sirich 2016). Studies indicate that indoxyl sulfate and *p*-cresyl sulfate promote the generation of reactive oxygen species, leading to oxidative stress and increased inflammatory cytokine production (Watanabe et al. 2013; Matsuo et al. 2015), which contribute to CKD pathogenesis and underscore the potential benefit of therapies targeting uremic toxin removal (Rossi et al. 2014; Sun, Chang, andWu 2012). Due to the limited removal of protein-bound uremic toxins such as indoxyl sulfate and *p*-cresol sulfate by conventional hemodialysis, oral activated carbon-based adsorbents like AST-120 have been developed to bind these toxins in the gut (Maheshwari et al. 2021; Lim et al. 2021). AST-120 is clinically used in Japan, Korea, the Philippines, and Taiwan, and has demonstrated the ability to lower circulating PBUTs (Asai, Kumakura, andKikuchi 2019). However, its efficacy in slowing CKD progression remains under investigation (Chen et al. 2019; Su et al. 2021), and its use is often limited by gastrointestinal side effects, including constipation (Akizawa et al. 2009; Schulman et al. 2006). These limitations highlight the need for alternative or complementary approaches. Biotherapeutic products, containing live microorganisms, may offer a less invasive strategy to reduce PBUTs (Cordaillat-Simmons, Rouanet, and Pot 2020). Synbiotics, which combine probiotics and prebiotics, have shown beneficial effects on gut health (Swanson et al. 2020). For example, in a vitro study, 33 *Bifidobacterium* and 26 *Lactobacillaceae* strains were pre-cultured for 16 hours, then incubated with 1 mM indole and *p*-cresol for 48 hours at 37°C, resulting in average removals of 32.4% indole and 4.2% *p*-cresol by *Bifidobacterium* and 7.5% indole and 17.6% *p*-cresol by *Lactobacillaceae*, demonstrating the potential of probiotics to reduce these uremic toxins (Candeliere et al. 2022). The in vivo study further demonstrated that this symbiotic treatment lowered serum indoxyl sulfate levels, improved estimated glomerular filtration rate, and alleviated gastrointestinal symptoms, highlighting its potential benefit for patients with kidney disease (Mitrović et al. 2023).

The main challenge in developing an effective biotherapeutic product for CKD patients is identifying active microorganisms that can remove protein-bound uremic toxins, such as indole and *p*-cresol, while functioning effectively in the colon. An optimized microbial delivery system can help overcome this challenge by protecting the microorganisms from harsh environmental conditions, such as low pH in stomach, encountered during passage through the digestive tract. Cell encapsulation in hydrogel have been developed to enhance the survival and efficacy of probiotics (Dafe et al. 2017; Systems 2018). Hydrogel coating layers protect microorganisms within the matrix, enabling them to withstand adverse conditions in the digestive tract (Li and Zhang 2024). Encapsulation also retains microbial biomass during the 20–40 hour colon transit, preventing cells from being washed out (LP and SF 1996). Furthermore, hydrogel encapsulation increases local microbial concentration, thereby enhancing catabolic activity (Minh et al. 2021). Cell densification within the hydrogel reduces the volume required for ingestion, which is especially important for CKD patients, who often must adhere to strict fluid intake restrictions (Wagner et al. 2022).

Although conventional probiotics like *Bifidobacterium* and *Lactobacillus* survive colon transit, their ability to remove protein-bound uremic toxins, such as indole and *p*-cresol, is limited, emphasizing the need for more potent strains. *Thauera aminoaromatica* S2 has shown high potential for *p*-cresol removal, and our previous study demonstrated its efficacy when encapsulated and delivered in a hydrogel under gut-like conditions (Saingam et al. 2025). In the present study, we aim to further evaluate the encapsulated *T. aminoaromatica* S2’s ability to remove p-cresol with co-presence of another protein-bound uremic toxin, indole. The study investigated (i) *T. aminoaromatica* S2 utilization of and tolerance to p-cresol and indole, (ii) *p*-cresol removal by encapsulated *T. aminoaromatica* S2 with co-presence of indole, and (iii) alleviating effect of incorporated activated carbon on p-cresol removal by encapsulated *T. aminoaromatica* S2 with co-presence of indole. Figure 1 depicts the concept of using hydrogel beads encapsulating *Thauera aminoaromatica* S2 to degrade *p*-cresol and indole in the intestine. This prevents these metabolites from entering the bloodstream and being converted in the liver into PBUTs, which are poorly cleared by dialysis.

**Figure 1.**
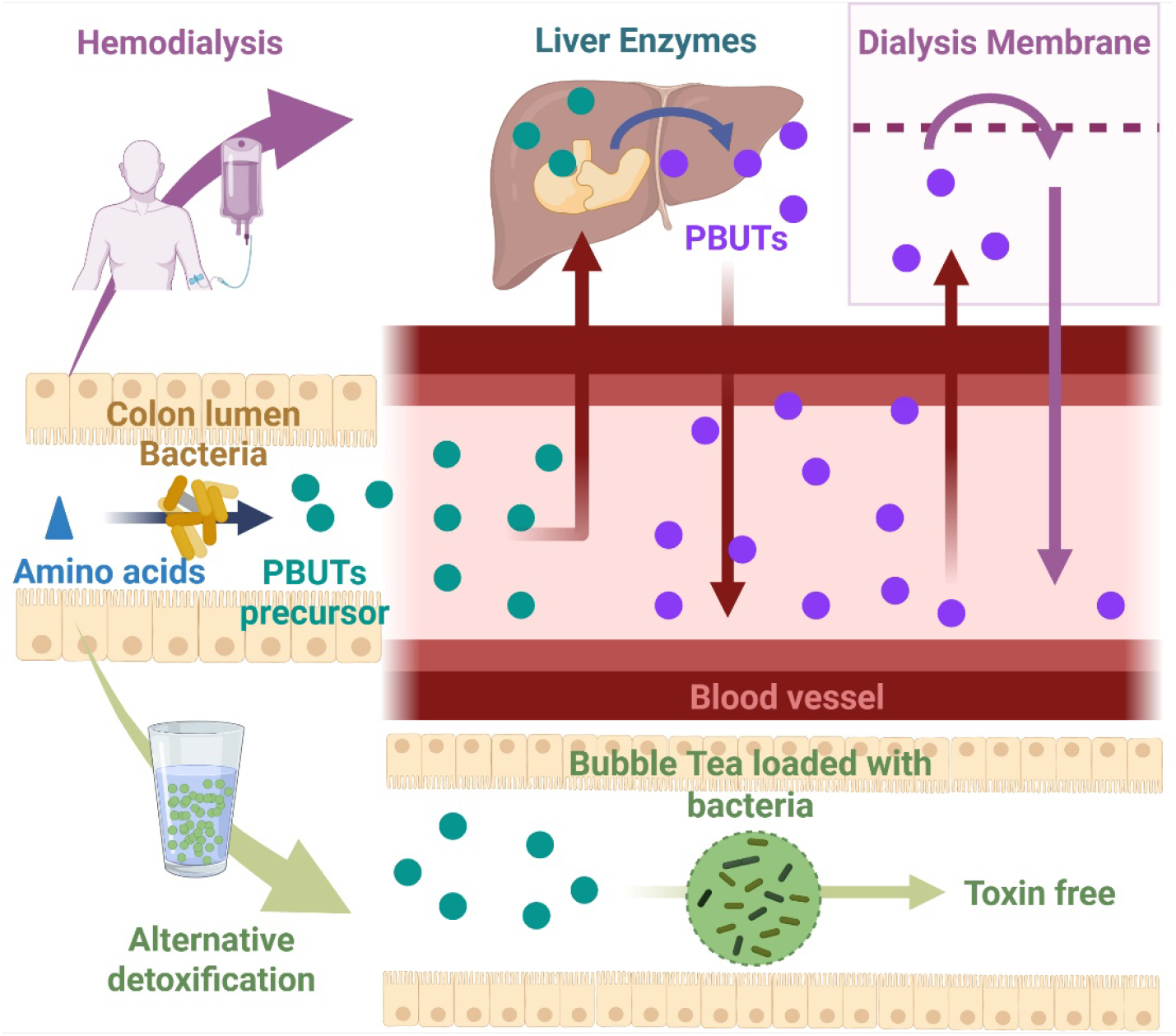
Protein-bound uremic toxins (PBUTs) in patients with CKD are not effectively removed by dialysis, whereas alternative gut-based detoxification can prevent their formation by eliminating precursors before absorption. Amino acids (dietary tryptophan and tyrosine) are converted in the intestine to indole and *p*-cresol, respectively. The bottom of the figure shows these metabolites being removed by bacteria encapsulated in hydrogel beads, preventing their entry into the bloodstream. The top of the figure illustrates the absence of this detoxification, where indole and *p*-cresol diffuse into the blood and are subsequently converted into PBUTs, which accumulate because dialysis cannot clear them. Figure created with [BioRender](https://www.biorender.com/).

## Material and method

### Bacteria culture and biomass cultivation

*T. aminoaromatica* S2 (DSMZ 14742) was obtained from the DSMZ collection (Braunschweig, Germany). The culture purity was confirmed and the liquid subcultures were stored in glycerol stocks at -80°C. In order to obtain the biomass for each experiment, *T. aminoaromatica* S2 was revived from glycerol stocks by aerobically plating on R2A media at 30°C for 48 h. A single colony was used as an inoculum for making the liquid subcultures. The bacteria cultivation media used for preparing the liquid subcultures was a basal media supplemented with vitamin solution, mineral solution, NaH_2_PO_4_, NaHCO_3_, Na_2_CO_3_, with 30 mM acetate as a carbon source and 25 mM NO_3_-as an electron acceptor under anaerobic cultivation at 30°C (Saingam et al. 2025). The bacteria cell pellet was collected by centrifuging the liquid subculture at 3,214 ×g for 25 min, and washed twice with the basal media. The washed cells were used for each experiment.

### Hydrogel encapsulation of *T. aminoaromatica* S2

The washed bacteria cell was resuspended in 6% w/v polyvinyl alcohol (PVA; Mw = 89,000– 98,000, 99+% hydrolyzed, Sigma-Aldrich, St. Louis, MO) and 2% sodium alginate (SA; Cape Crystal Brands, Summit, NJ) solution with cell encapsulation density of ∼5 Log CFU (mL hydrogel)^*-*1^. The hydrogels were made by extruding the cell resuspension through a 21G sterilized syringe needle into 4% (w/v) CaCl_2_, which were stirred for 1 h in the anaerobic atmosphere. Anaerobic conditions for the bacteria cultivation and hydrogel encapsulation was achieved using Hungate techniques (Hungate, 1969) or using the Coy(TM) anaerobic chamber (85% N_2_: 10% CO_2_: 5% H_2_ atmosphere).

### Indole effect on *T. aminoaromatica* S2 growth

The washed bacteria cells were resuspended in the basal media and the cell resuspension was added to 96 well-plates with 10 mM NO_3_-as an electron acceptor and varied test conditions of *p*-cresol (0, 0.5, 1, 2, 4, and 8 mM) and/or indole (0, 0.25, 0.5, 1, 2, 4, and 8 mM). Each condition was performed with three replicate wells. Initial bacteria cell number in each well was approximately ∼10^6^. The plates were incubated in resealable pouches with anerobic atmosphere-generating GasPak EZ Anaerobe Container System Sachet (Becton Dickinson, Franklin Lakes, New Jersey) at 37°C for three days. The OD_600_ of the test resuspensions was read prior to and after three-day incubation to measure the bacteria growth. The wells without carbon sources were included and the arithmetic average of their OD_600_ readings were used as the threshold to determine bacteria growth in other wells.

### Indole effect on *p*-cresol removal by encapsulated *T. aminoaromatica* S2

Fifteen aliquots of ∼one mL of the bacteria hydrogels were prepared and experimented. Each hydrogel aliquot was transferred to a Balch-type tube containing 12 mL of the bacteria cultivation media modified for the hydrogel test as mentioned in our previous study (Saingam et al. 2025), supplemented with 2 mM *p*-cresol and varied concentrations of indole (0, 0.25, 0.5, and 1 mM). Each experiment condition was performed using three aliquots of the bacteria hydrogels. The other three aliquots of the bacteria hydrogels, used as control, were prepared using heat-disinfected bacteria cells and incubated in the media supplemented with 2 mM *p*-cresol and 1 mM indole. The tubes were incubated at 37°C in a shaker at 133 rpm. At 0, 12, 24, 42, 66, 88, 112 h, liquid samples were collected using Hungate techniques and processed for the chemical analysis.

### Indole effect on *p*-cresol removal by encapsulated *T. aminoaromatica* S2 with activated carbon

Nine aliquots of ∼one mL of the bacteria hydrogels were prepared and experimented. The 18G needle was used for extruding hydrogel suspension due to activated carbon particle sizes. Each hydrogel aliquot was transferred to a Balch-type tube containing 12 mL of the bacteria cultivation media modified for the hydrogel test as mentioned in our previous study (Saingam et al. 2025), supplemented with 2 mM *p*-cresol and 1 mM indole. Among the nine aliquots of the bacteria hydrogels, three aliquots were live bacteria cells, and three aliquots were live bacteria cells with sterile activated carbon (Fisher Chemical, Waltham, Massachusetts) of ∼0.1 g/ mL hydrogel density. The other three aliquots, which were heat-disinfected bacteria, were used as control. In addition, the other three aliquots of sterile activated carbon hydrogels (∼0.1 g/ mL hydrogel density) were prepared as the activated carbon control. The tubes were incubated at 37°C in a shaker at 133 rpm. At 0, 13, 17, 25, 40, and 64 h, liquid samples were collected using Hungate techniques and processed for the chemical analysis.

### Chemical analysis

The liquid samples were filtered using 0.2 µm nylon filter (VWR, Radnor, PA) prior to measurements of *p*-cresol, indole, total nitrogen (NO_3_- and NO_2_-), and NO_2_-. Infinity II liquid chromatography (LC) system (Agilent Technologies, CA) with Aminex HPX-87H column (Biorad laboratories, CA) was used to measure *p*-cresol and indole concentrations. The mobile phase was acetonitrile and 0.5 mM H_2_SO_4_ (2:8 volume ratio) with the flow rate of 0.6 mL/min and the column temperature of 30°C. The chromatogram signals were measured at UV 210 nm. Gallery^TM^ Automated Photometric Analyzer (Thermo Fisher Scientific, Waltham, MA) was used to measure total nitrogen and NO_2_-spectrophotometrically using the reagents and following manufacturer instructions. The absolute p-cresol and indole removal rates were calculated using the measurements of the initial and final samplings. The removal rates were reported as nmol (mL hydrogel)^-1^ h^-1^. The arithmetic mean and standard deviations were calculated from the measurements of three biological replicates. Comparison of removal rates between two different conditions were achieved using Mann-Whitney U Test. Significance level was at 0.05.

### Bacteria cell quantification

Hydrogels and liquid samples were collected for bacteria cell quantification through the plate counting method and/or the flow cytometry. The collected hydrogels were rinsed with sterile water and dissolved in 1 mL of 20 mM sodium citrate in phosphate buffer solution (PBS) (Saingam et al. 2025). The hydrogel suspension was incubated at room temperature until complete dissolution of the hydrogels. Serial dilutions of the dissolved hydrogel suspension and collected liquid sample were obtained using PBS and plated on R2A media. The plates were aerobically incubated at 30 °C for 5 days and only the plates with observed colony numbers of 30-300 were used to determine bacteria cell quantification. The dissolved hydrogel suspension and collected liquid sample were stained with SYBR in the dark condition for 30 min at 37 °C. The cells in stained suspension were quantified using the Guava EasyCyte flow cytometer (Luminex, Austin, USA). The arithmetic mean and standard deviations were calculated from the measurements of three biological replicates.

## Result

### Indole and *p*-cresol effect on *T. aminoaromatica* S2 growth

After three days of incubation with a 0.5-8.0 mM *p*-cresol or 0.25-8.0 mM indole as sole carbon source, *T. aminoaromatica* S2 was able to utilize *p*-cresol at concentrations of 0.5, 1, and 2 mM, and indole at 0.25 mM, as indicated by growth relative to the control that was grown without any carbon addition (Figure 2), confirming that both *p*-cresol and indole can serve as sole carbon sources for its metabolism. Growth was inhibited at higher concentrations, with indole exerting stronger inhibition at 0.5 mM, whereas *p*-cresol inhibition occurred only at 4 mM, demonstrating differential tolerance of *T. aminoaromatica* S2 to these uremic toxins. The presence of both p-cresol and indole carbon sources exhibited a synergistic effect on *T. aminoaromatica* S2 growth pattern compared to conditions with a single carbon source (Figure 3). While growing on 2 mM *p*-cresol, the addition of 0.25 mM indole enhanced growth compared to the corresponding *p*-cresol-only condition. In addition, the presence of *p*-cresol alleviated the inhibitory effect of higher indole concentrations; for example, 0.5 mM *p*-cresol allowed growth at 1 mM indole, whereas no growth occurred at 1 mM indole when it was the sole carbon source. These results indicate that *T. aminoaromatica* S2 can utilize *p*-cresol and indole concomitantly, with *p*-cresol mitigating indole-induced growth inhibition.

**Figure 2.**
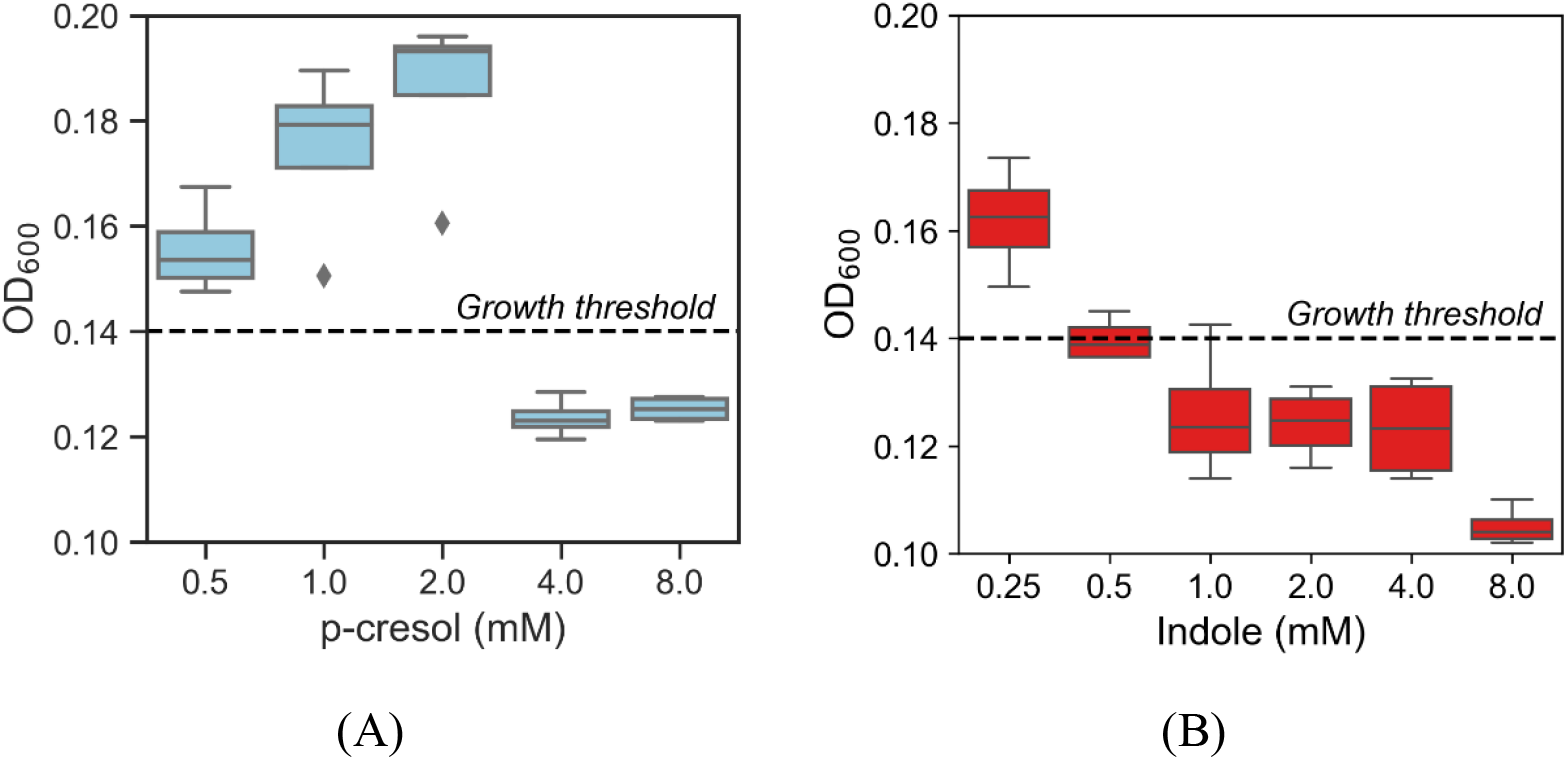
Growth of *T. aminoaromatica* S2 fed with (A) 0.5 – 8.0 mM *p*-cresol and (B) 0.25 – 8.0 mM indole. The boxplots represent mean and data distribution across four replicates.

**Figure 3.**
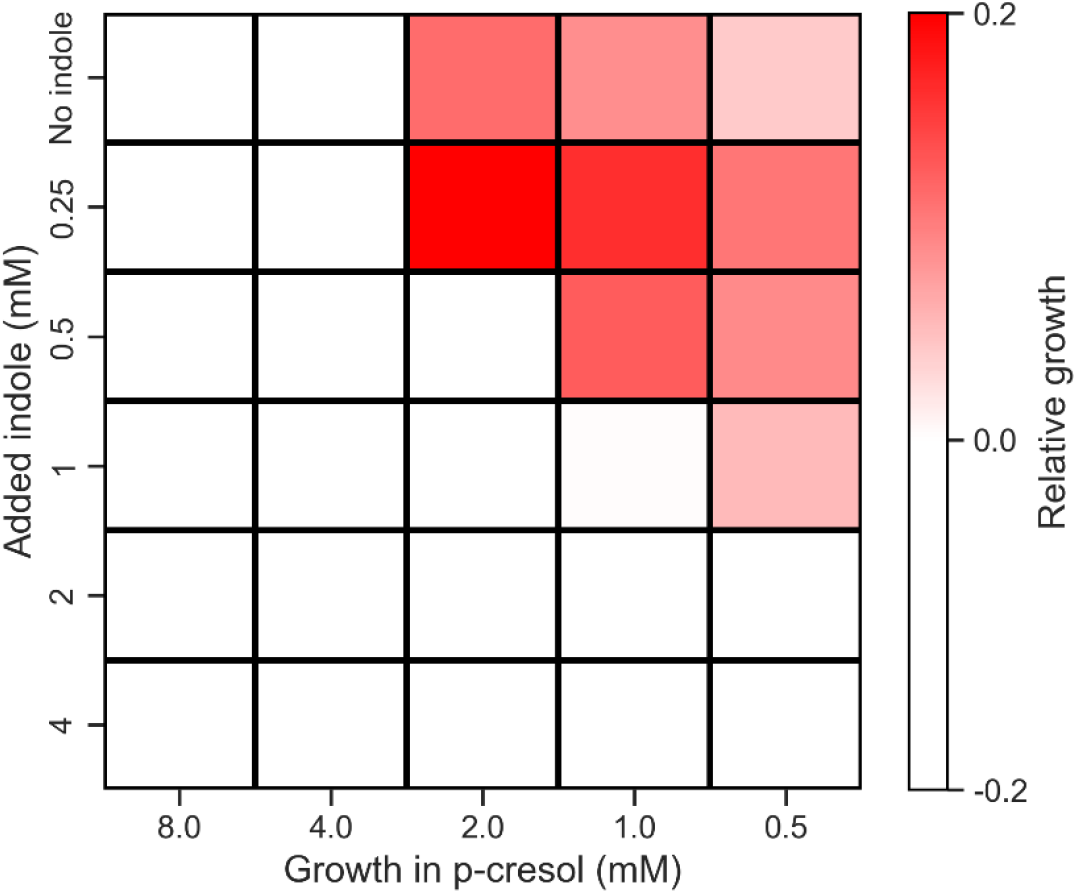
Effect of indole on growth of *T. aminoaromatica* S2 fed with 0.5 – 8.0 mM *p*-cresol. The added concentrations of indole was 0 – 4 mM. The relative abundance represented logarithm of the ratio of the OD_600_ average of the four replicates of test cultures to the OD_600_ average of the control culture.

### Indole effect on *p*-cresol removal by encapsulated *T. aminoaromatica* S2

*T. aminoaromatica* S2 were encapsulated in PVA/SA hydrogels and tested for removal of 2 mM *p*-cresol with co-presence of varied indole concentrations (0 - 1 mM) (Figure 4). When there was no co-presence of indole, the hydrogels of *T. aminoaromatica* S2 showed the highest *p*-cresol removal (Figure 4A). In contrast to growth pattern observed with planktonic cells, the hydrogels of *T. aminoaromatica* S2 exposed to 0.5 mM indole showed p-cresol absolute removal rate similar to those exposed to 0.25 mM (Figure 4A). The 1 mM indole showed inhibitive effect on the *p*-cresol removal of encapsulated *T. aminoaromatica* S2 as indicated by the absolute removal rate similar to the control hydrogels of heat-disinfected cells (Figure 4A). During the test, the Balch tubes of the control hydrogels of heat-disinfected cells showed indole decrease due to physical absorption. The live encapsulated *T. aminoaromatica* S2 hydrogels with co-presence with 1 mM indole showed similar level of indole decrease.

**Figure 4.**
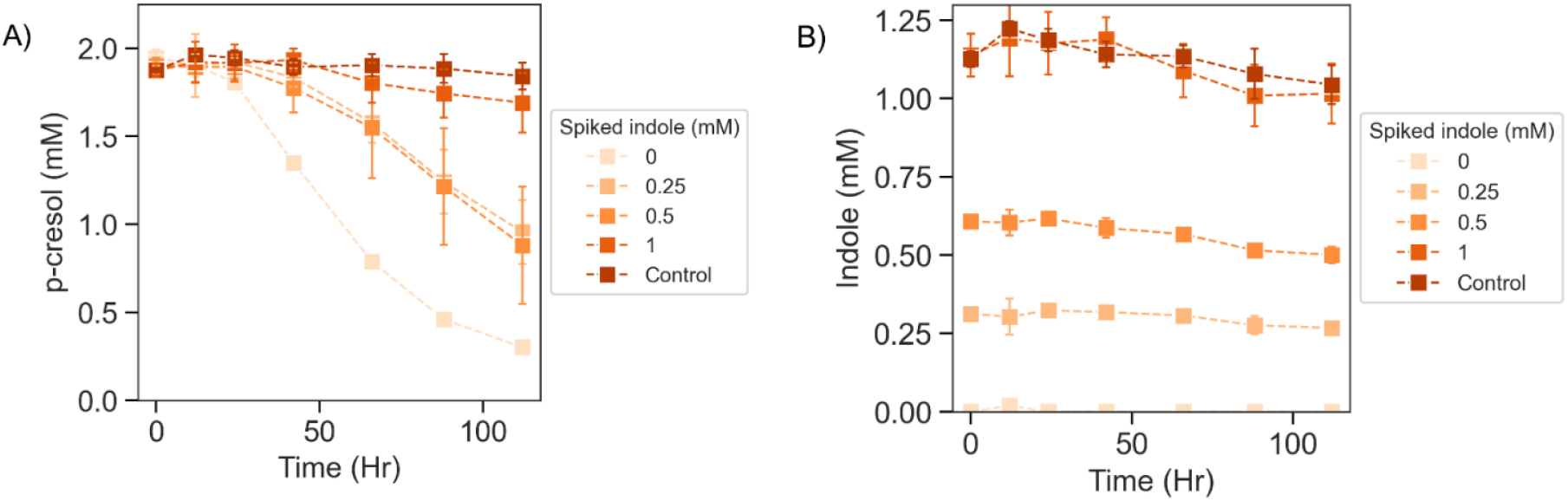
Effect of indole on p-cresol removal by encapsulated *T. aminoaromatica* S2. (A) p-cresol concentrations throughout the removal test and (B) residuals concentrations of spiked indole. The data showed average and standard deviations of three replicates.

### Indole effect on *p*-cresol removal by encapsulated *T. aminoaromatica* S2 with activated carbon

The addition of activated carbon enabled the removal of 2 mM *p*-cresol and 1.25 mM indole within 20 hours (Figure 5), demonstrating the benefit of activated carbon, amending hydrogels for enhanced *p*-cresol removal and the mitigation of indole toxicity.

**Figure 5.**
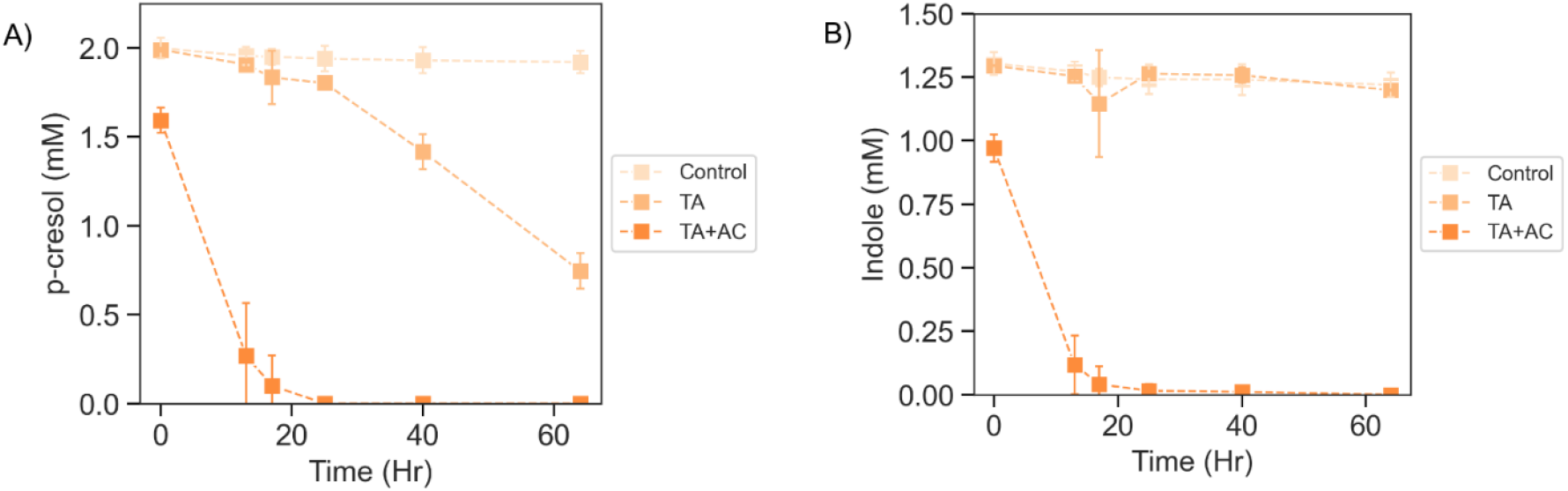
Improved *p*-cresol removal by encapsulated *T. aminoaromatica* S2 (TA) with activated carbon (AC). (A) *p*-cresol concentrations throughout the removal test and (B) residual concentrations of spiked indole. The data showed average and standard deviations of three replicates.

## Discussion

*T. aminoaromatica* S2 can utilize *p*-cresol but shows minimal growth on indole as a carbon source, consistent with Mechichi et al (Mechichi et al. 2002). In feces, indole and *p*-cresol concentrations range from 0.30–6.64 mM (Darkoh et al. 2015), while *p*-cresol concentrations have been reported to range from 0.6–5.2 mM (Saingam et al. 2025), based on previously published studies. When tested as the sole carbon source, the maximum concentrations supporting growth of *T. aminoaromatica* S2 were 0.5 mM for indole and 2 mM for *p*-cresol. Notably, the addition of 0.25 mM indole enhanced growth compared to the *p*-cresol-only condition, suggesting a potential synergistic effect. Consistently, *T. aminoaromatica* S2 was able to grow in the presence of 1 mM indole combined with 0.5 mM *p*-cresol, whereas no growth was observed with 1 mM indole as the sole carbon source. Together, these findings suggest that *T. aminoaromatica* S2 can maintain the *p*-cresol-degrading activity under co-exposure conditions relevant to the intestinal lumen, provided that local toxin concentration remain within a tolerable range.

Hydrogel encapsulation provides an effective strategy to increase local bacterial density, retain cells within the matrix, and enhance *p*-cresol removal, while also potentially improving the tolerance of *T. aminoaromatica* S2 to indole toxicity. In our experiments, when hydrogel-encapsulated cells were exposed to 2 mM *p*-cresol and 0.5 mM indole, no growth was observed in planktonic culture; however, *p*-cresol degradation still occurred in the encapsulated system. Although *T. aminoaromatica* S2 exhibited only slow and limited p-cresol removal at 2 mM *p*-cresol and 1 mM indole, hydrogel encapsulation still conferred greater tolerance to indole compared with planktonic cultures. The hydrogel matrix likely prevents cell washout and retains biomass, thereby maintaining sufficient cell density to support *p*-cresol and indole removal, which may be particularly beneficial for gut applications where elevated indole concentrations and continuous intestinal flow could otherwise limit bacterial persistence and activity. Importantly, these data support the broader concept that hydrogel delivery can enable deployment of non-native (environmental) strains with specialized metabolic capabilities that are not robustly expressed by the resident gut community, thereby enabling gut-localized removal of *p*-cresol at its source prior to absorption into the blood stream and downstream PBUT formation. While the present work is *in vitro*, the results provide a strong rationale for further development of encapsulated microbial “metabolic add-on” strategies as an oral approach to intercept uremic toxin precursors upstream of systemic exposure.

Interestingly, in planktonic cultures, *T. aminoaromatica* S2 showed maximal growth with 2 mM *p*-cresol and 0.25 mM indole compared with 2 mM *p*-cresol alone. In contrast, in hydrogel-encapsulated cultures, *p*-cresol removal was reduced in the presence of 0.25 mM indole relative to hydrogel batches containing only 2 mM *p*-cresol, indicating that hydrogel encapsulation alters the effect of indole on bacterial activity. This divergence highlights that performance in encapsulated formats cannot be inferred directly from planktonic behavior, because the hydrogel microenvironment changes mass transport, local substrate/toxin accumulation, and the effective exposure experienced by cells.

One possible explanation is that the hydrogel limits diffusion of *p*-cresol and indole to the cells, allowing *T. aminoaromatica* S2 to tolerate higher indole concentrations. In addition, the higher local cell density within the hydrogel may provide collective protection against toxicity. Conversely, the hydrogel may also retain indole or toxic metabolic intermediates, increasing local concentrations and thereby reducing bacterial activity, which could explain the lower performance compared with planktonic cultures under certain conditions.

The addition of activated carbon proved to be an effective strategy for removing both *p*-cresol (2 mM) and indole (1.25 mM) within 20 hours (reflective of typical gut transit times), whereas under the same conditions *T. aminoaromatica* S2 alone achieved approximately 75% *p*-cresol removal only after 60 hours. Despite the strong adsorption capacity of activated carbon for toxin removal, it has been reported to have the potential of binding essential nutrients (Ivanov et al. 2012), which could be detrimental for future clinical applications where maintaining nutrient availability for kidney patients is critical. Therefore, future studies could investigate reducing the amount of activated carbon in the hydrogel to selectively mitigate indole toxicity while preserving sufficient nutrients to support the activity of *T. aminoaromatica* S2. More broadly, these results suggest that combining encapsulated microbes with localized adsorption offers a modular strategy to stabilize microbial function in complex gut-relevant chemical environments, while maintaining a gut-centered approach for toxin removal at its source.

## Conclusion

Planktonic *Thauera aminoaromatica* S2 was able to utilize 2 mM *p*-cresol as a carbon source but showed growth only at relatively low indole concentrations (≤0.5 mM). A synergistic effect was observed during co-exposure to *p*-cresol and indole. Specifically, higher growth was observed with 0.25 mM indole plus 2 mM *p*-cresol compared with 2 mM *p*-cresol alone, and growth was supported with 1 mM indole plus 0.5 mM *p*-cresol, whereas no growth occurred with 1 mM indole in the absence of *p*-cresol. These findings indicate the potential of *Thauera aminoaromatica* S2 to remove *p*-cresol in the gut environment, where both indole and *p*-cresol are present.

Hydrogel encapsulation further expanded the tolerance of *Thauera aminoaromatica* S2 to indole toxicity, increasing the maximum tolerated indole concentration from 0.25 mM to 1 mM while coexisting with 2 mM *p*-cresol. Encapsulated cells initiated *p*-cresol removal after approximately 50 hours, although at a reduced rate compared with lower-toxicity conditions. The incorporation of activated carbon into the hydrogel further enhanced the removal of both indole and *p*-cresol and shortened the effective treatment time to better match colonic transit time, suggesting that encapsulated *Thauera aminoaromatica* S2 could be a promising future therapeutic approach for patients with chronic kidney disease. Collectively, these results support hydrogel encapsulation as a promising oral delivery format for environmental strains that provide metabolic functions not robustly present in the native gut microbiome, enabling interception of *p*-cresol in the intestinal lumen prior to systemic absorption. Further optimization of bead design and co-formulated additives may improve removal kinetics and robustness, supporting next-step evaluation of this strategy.

## AI disclosure

AI-assisted tools (OpenAI’s ChatGPT) were used to help rewrite sections of the manuscript and to draft the abstract. All scientific content, interpretations, and conclusions were reviewed and verified by the authors.

## Acknowledgments

This project was funded in part by NIH (1R01DK130815-01).

